# Genomic knockout of *alms1* in zebrafish recapitulates Alström syndrome and provides insight into metabolic phenotypes

**DOI:** 10.1101/439067

**Authors:** Jessica E. Nesmith, Timothy L. Hostelley, Carmen C. Leitch, Maggie S. Matern, Saumil Sethna, Rebecca McFarland, Sukanya Lodh, Christopher J. Westlake, Ronna Hertzano, Zubair M. Ahmed, Norann A. Zaghloul

## Abstract

Alström syndrome is an autosomal recessive obesity ciliopathy caused by loss-of-function mutations in the *ALMS1* gene. In addition to multi-organ dysfunction, such as cardiomyopathy, retinal degeneration, and renal dysfunction, the disorder is characterized by high rates of obesity, insulin resistance and early onset type 2 diabetes mellitus (T2DM). To investigate mechanisms linking disease phenotypes we generated a loss-of-function deletion of a*lms1* in the zebrafish using CRISPR/Cas9. We demonstrate conserved phenotypic effects including cardiac defects, retinal degeneration, and metabolic deficits that included propensity for obesity and fatty livers in addition to hyperinsulinemia and glucose response defects. Gene expression changes in β-cells isolated from *alms1^−/−^* mutants revealed changes consistent with insulin hyper-secretion and glucose sensing failure, which were also identified in cultured murine β-cells lacking *Alms1*. These data present a zebrafish model to assess etiology and new secretory pathway defects underlying Alström syndrome-associated metabolic phenotypes. Given the hyperinsulinemia and reduced glucose sensitivity in these animals we also propose the *alms1* loss-of-function mutant as a monogenic model for studying T2DM phenotypes.

**AUTHOR SUMMARY:** These data comprise a thorough characterization of a zebrafish model of Alström syndrome, a human obesity syndrome caused by loss-of-function deletions in a single gene, *ALMS1*. The high rates of obesity and insulin resistance found in these patients suggest this disorder as a single-gene model for Type 2 Diabetes Mellitus (T2DM), a disorder caused by a variety of environmental and genetic factors in the general population. We identify a propensity for obesity, excess lipid storage, loss of β-cells in islets, and hyperinsulinemia in larval and adult stages of zebrafish *alms1* mutants. We isolated β-cells from the *alms1* mutants and compared the gene expression profiles from RNASeq datasets to identify molecular pathways that may contribute to the loss of β-cells and hyperinsulinemia. The increase in genes implicated in generalized pancreatic secretion, insulin secretion, and glucose transport suggest potential β-cell exhaustion as a source of β-cell loss and excess larval insulin. We propose this mutant as a new genetic tool for understanding the metabolic failures found in Type 2 Diabetes Mellitus.

## INTRODUCTION

Primary cilia are present on the majority of vertebrate cell types and act as cellular hubs for sensing and transducing signaling pathways (1). Deletions in ciliary genes—including basal body proteins, transition zone components, and intraflagellar transport elements—are implicated in a class of diseases termed the ciliopathies which exhibit a broad range of phenotypes (2,3). A unique subset among these disorders are the obesity ciliopathies that present with highly penetrant, early onset obesity. Alström syndrome, one of two major obesity ciliopathies, is associated with the most profound metabolic derangement among the disorders, including prominent truncal obesity, severe insulin resistance, hyperinsulinemia, early onset Type 2 Diabetes Mellitus (T2DM), and several other metabolic syndrome features (4,5). This rare autosomal recessive syndrome is caused by pathogenic variants in *ALMS1*, which localizes to the basal body of cilia (6), and may play a role in intracellular trafficking (7,8). Additional hallmark features of the disorder include cardiomyopathy, retinal degeneration, and renal dysfunction, as well as childhood obesity and T2DM. The unique features of Alström syndrome, however, offer the opportunity for generating models to better understand the cellular etiology underlying metabolic conditions, including T2DM, and to investigate the potential role of ciliary proteins.

Here, we present a zebrafish model of Alström syndrome. Zebrafish (*Danio rerio*) have long been used as models of human genetic disease. The external embryonic development allows for whole-body knockdown and knockout studies, the wide range of existing transgenic reporters and translucent body plan permits high-resolution microscopy of whole animals, and the high fecundity provides ease of mutant generation as well as large scale -omic studies (9–12). Zebrafish have also recently emerged as robust models of metabolic traits owing to conservation of basic physiology, including gastrointestinal function, endocrine regulation, lipid and glucose metabolism, and central neuronal regulation of food intake, all present by 5 days post-fertilization(dpf) (13,14). We previously demonstrated a link between β-cell proliferation and obesity ciliopathies using knockdown models of two ciliopathies, Bardet-Biedl syndrome (BBS) and Alström syndrome (8,15). The latter was carried out by transient knockdown of the zebrafish *alms1* gene during early embryonic stages. While these studies inform the role of *alms1* in regulation of β-cell production early in life, the inherent limitations of transient knockdown preclude more extensive investigation of other later onset and adult phenotypes.

In the present study, we generated a genomic model of Alström syndrome by targeting the zebrafish *alms1* gene using CRISPR/Cas9. The resulting heritable mutation ablated protein production and resulted in systemic defects that recapitulate the human syndrome and existing mouse models, including defects in neurosensory, cardiac, renal, and metabolic systems. We further explored metabolic phenotypes and found highly penetrant hepatic steatosis, increased weight gain under high fat feeding conditions, impaired glucose uptake, defective β-cell response to high-glucose conditions, and early onset hyperinsulinemia. To more closely examine glucose regulation in Alström syndrome, we introduced the loss-of-function mutation in *alms1* into a transgenic β-cell reporter line. We used this line to selectively isolate β-cells from the *alms1^−/−^* zebrafish and examine whole transcriptome data using RNA-seq. These data support a role for *alms1* in modulation of β-cell insulin secretion through regulation of intracellular secretion and also provide evidence of impaired β-cell glucose-sensing. These findings indicate β-cell autonomous defects are a primary driver of early onset hyperinsulinemia and early onset T2DM, providing insight into β-cell defects underlying more common forms of T2DM.

## RESULTS

### Generation of a zebrafish *alms1* genomic loss-of-function mutant

We targeted zebrafish *alms1* with CRISPR/Cas9 by injection of guide RNAs (gRNA) targeted to exon 4 of the gene along with Cas9 protein injection directly into 1-cell stage embryos (Fig 1A). To allow for both whole body phenotyping as well as β-cell-specific defects in mutants, we deleted *alms1* in the Tg(*insa:mCherry*) transgenic zebrafish in which β-cells express the mCherry fluorophore under control of the *preproinsulin (insa)* promoter (16). In adult F1 animals, we identified a heritable 7 bp deletion (c.1086_1092del) in exon 4 that is predicted to result in a frameshift and introduction of a premature stop codon at p.S364X (Fig 1A). We identified multiple heterozygous F2 carriers of the deletion and in-crossed them to propagate a homozygous *alms1^−/−^* line. We noted non-Mendelian ratios in progeny of heterozygous parents, with homozygotes representing only 14% of genotyped adult fish (p=0.007; Chi-Squared; Fig 1B), suggesting an important developmental role for *alms1*. *alms1^−/−^* homozygous mutant animals survived to adulthood and could be mated to produce viable progeny. Their offspring also exhibited high rates of curly tail phenotype at 3 days post-fertilization (dpf), which is typical of zebrafish ciliary mutants (Fig 1C) (17). *alms1* RNA and Alms1 protein levels were dramatically decreased in *alms1^−/−^* animals (Fig 1D-E). Intermediate RNA levels were identified in heterozygote animals, although no morphological abnormalities were observed (Fig S1A). Interestingly, visible morphological defects including the body curvature phenotype were rarely observed in homozygous progeny of *alms1^+/−^* parents, potentially suggesting a maternal contribution of *alms1* to early development.

**Fig 1.**
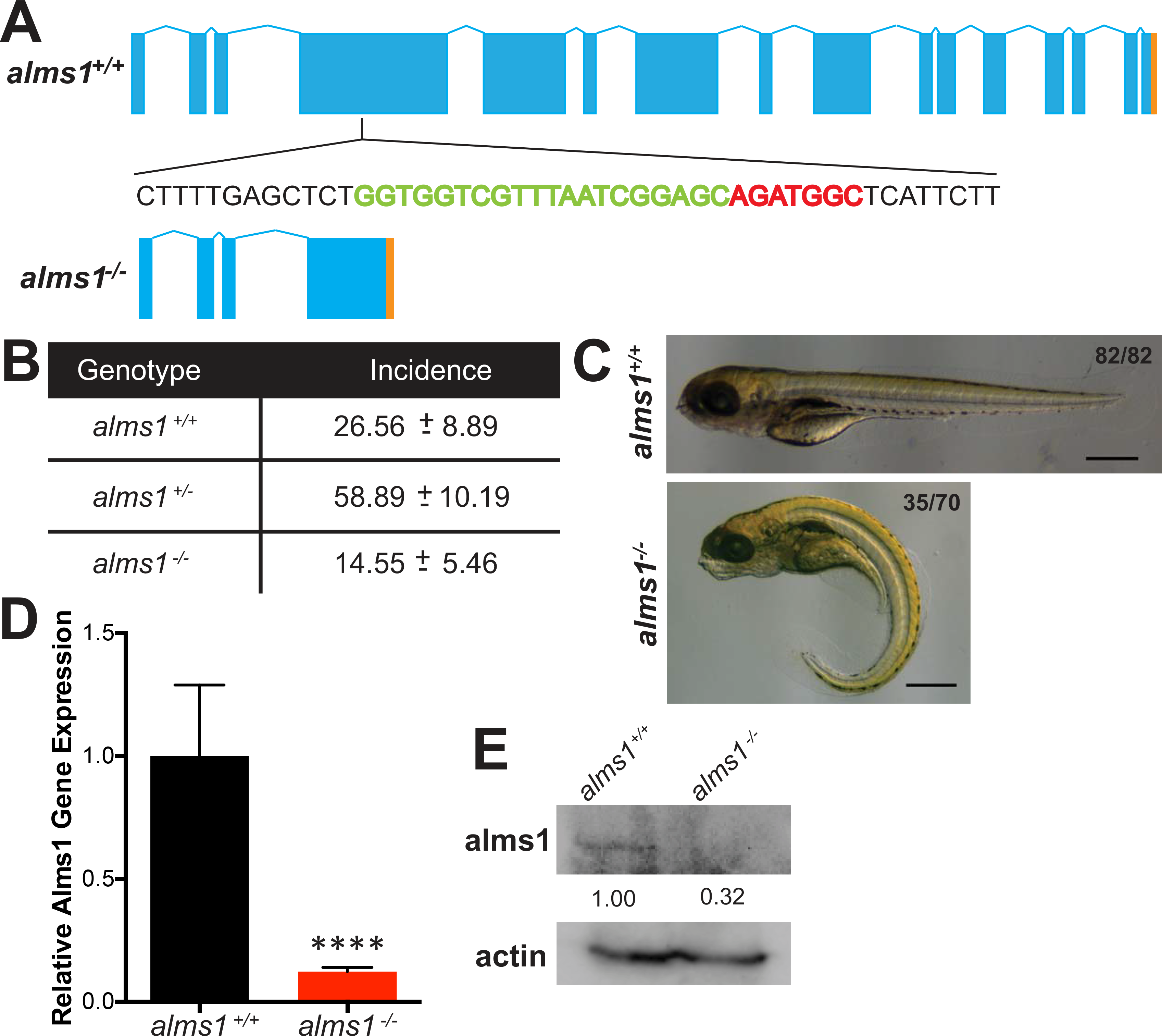
Generation of zebrafish *alms1^−/−^* line. (A) Schematic of alms1 genomic region with and without CRISPR/Cas9 induced deletion resulting in premature stop codon (orange). Pull out indicates region in exon 4 with sgRNA target (green) and deletion (red) generated using CRISPR/Cas9. (B) Ratios of identified mutants in heterozygous in-cross progeny. Note the reduced rate of mutants relative to expected mendelian ratios. (C) Representative images of *alms1^+/+^* (n=82) and *alms1^−/−^* (n=70) larvae at 5 days post-fertilization. Note the ciliopathy body dysmorphogenesis identified in *alms1^−/−^* larvae. Scale bar, 1 mm. (D) *alms1^−/−^* larvae demonstrate reduced cDNA expression levels compared to *alms1^+/+^* at 5 days post-fertilization (dpf). Statistics, Student’s t-test, ****p<0.0001. (E) *alms1^−/−^* larvae demonstrate reduced protein levels compared to *alms1^+/+^* at 5 dpf.

### Zebrafish *alms1^−/−^* phenocopies features of Alström syndrome including cardiomyopathy, retinal dystrophy and renal defects

Having identified viable homozygous *alms1* mutants, we examined broader phenotypes in *alms1^−/−^* progeny. Alström syndrome patients and mouse mutant models exhibit a wide range of phenotypes across multiple organ systems (4,18). We examined mutants to assess the presence of prominent features associated with the disorder in both embryonic and adult tissues.

Nearly two-thirds of Alström syndrome patients present with dilated cardiomyopathy and congestive heart failure contributes to a significant portion of the reported mortality (19). The earliest manifestation of cardiomyopathy in zebrafish is cardiac edema, which results from failed contractility, loss of vascular integrity or ventricular malformations and can develop by 48 hpf (20). Compared to 2.5% of wild type control animals generated from sibling clutch-mates, 18% of *alms1^−/−^* embryos exhibited cardiac edema at 48 hpf (Fig 2A-B). Although this reflected a 6-fold increase in edema (p<0.0001; Observed vs Expected), the majority of *alms1^−/−^* embryos appeared largely normal (Fig 2A-B). To examine the possibility of a progressive cardiac phenotype, we also examined the morphology of age-matched adult hearts in *alms1^−/−^* animals. We observed smaller hearts in the adult *alms1^−/−^* fish when compared with wildtype controls (Fig 2C’). Closer examination revealed a loss of ventricular wall integrity in the *alms1^−/−^* fish (Fig 2C”). These findings are consistent with cardiomyopathy in Alström, including the reported variability in phenotypic presentation in patients (19).

**Fig 2.**
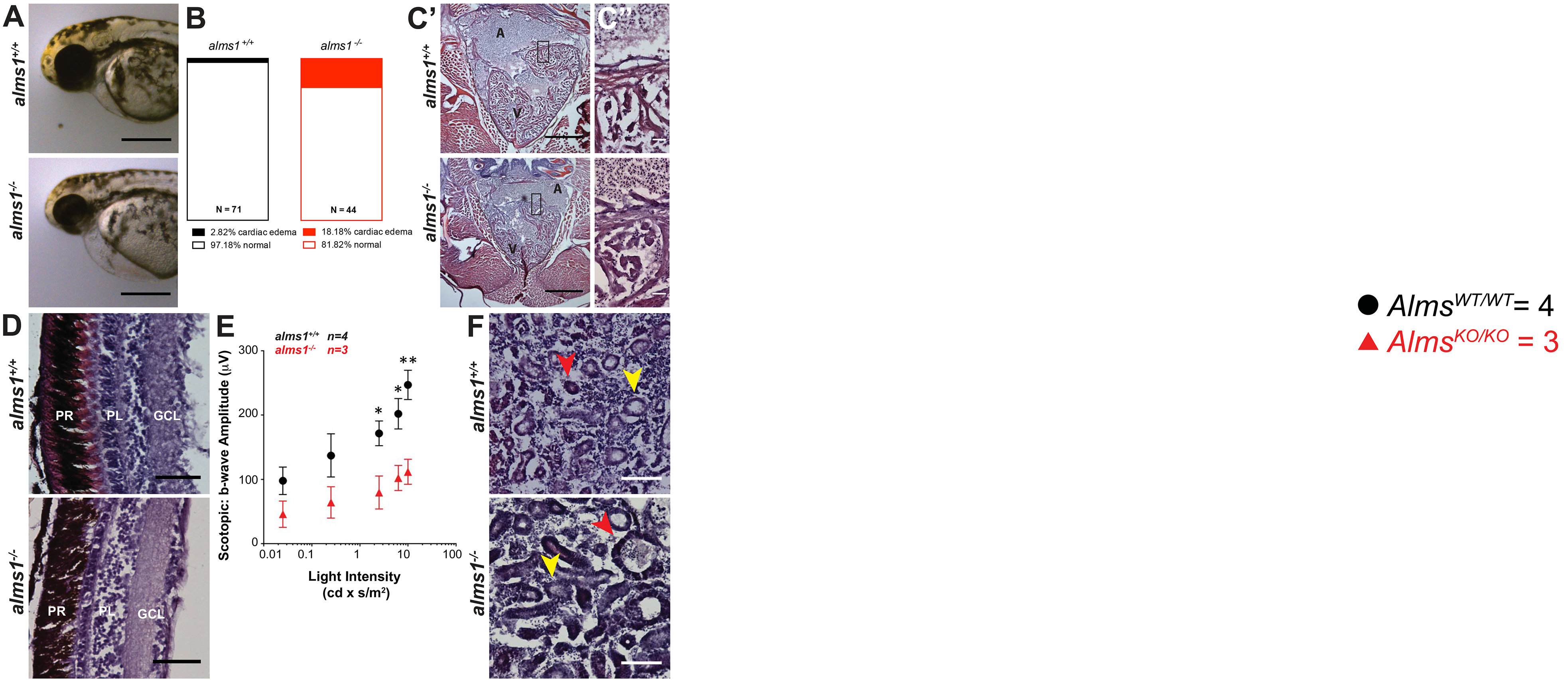
*alms1^−/−^* zebrafish display multiple defective organ systems. (A) Representative images of *alms1^+/+^* and *alms1^−/−^* larvae showing severe cardiac edema at 48 hours post-fertilization. Scale bar, 1 mm. (B) Quantification of cardiac edema rates in *alms1^+/+^* (n=71) and *alms1^−/−^* (n=44) larvae. Significance, Chi-squared, **** p<0.0001. (C) Adult cardiac H#E sections from *alms1^+/+^* and *alms1^−/−^* zebrafish at 6 months showing gross (scale bar, 500 μm; C’) and high-magnification (scale bar, 25 μm; C’’) structures in the atrium and ventricle of the heart. Note the diminished size and degraded ventricular wall composition in the *alms1^−/−^* animals. A: atrium, V: ventricle. (D) Representative images of H#E staining of retinal layers. Note degradation of multiple retinal layers in alms1^−/−^ animals. PR: photoreceptors, PL: plexiform layers, GCL: ganglion cell layers. Scale bar, 50 μm. (E) Quantification of scotopic b-wave amplitudes in response to light stimulation from *alms1^+/+^* (n=4) and *alms^−/−^* (n=3) zebrafish at 9 months of age. (F) Representative H#E images of kidneys in *alms1^+/+^* and *alms1^−/−^* animals. Note the abnormal shape and size of kidney tubules in *alms1^−/−^* animals. Distal tubule, yellow arrow, proximal tubule, red arrow. Scale bar,100 μm. Where indicated, *p<0.05, **, p<0.01.

Degeneration of the retina is also typical in Alström patients with individuals presenting with visual impairment within a few weeks of birth and typically becoming completely blind before age 30 as a result of progressive loss of photoreceptors (4,21). We assessed retinas of *alms1^+/+^* adults and found no impairment of the retinal architecture, with the ganglion cell layer, the plexiform layers, and photoreceptor layers all intact (Fig 2D). The *alms1^−/−^* adults exhibited reduced retinal integrity, most notably in both the inner and outer photoreceptor layers which were visibly reduced (Fig 2D). These observations are consistent with the retinal degeneration and subsequent blindness found in human patients. To assess functional impact of *alms^−/−^* on vision, we performed full-field electroretinograms (ERGs) on *lms1^+/+^* and *alms^−/−^* zebrafish at 9 months of age. Non-invasive *in vivo* scotopic ERGs showed attenuated b-wave amplitudes in the *alms^−/−^* zebrafish as compared to controls (Fig 2E), indicating reduced photoreceptor response in these animals. Intriguingly, we noticed age-dependent reduced motility and decreased food seeking behavior in *alms1^−/−^* mutants, consistent with severe vision loss (data not shown). Together with the histological findings, these data implicate Alms1 in maintenance of retinal sensory epithelia and visual function.

Nearly half of Alström patients have reported renal failure accompanied by a broad range of defects including calculi, interstitial fibrosis, and glomerulosclerosis (4). We examined gross kidney morphology in H#E-stained sections of kidneys from *alms1^+/+^* adults, and observed well-formed tubule structure of varying intensities, indicating distal (lighter stain, yellow arrow) and proximal (darker stain, red arrow) tubules (Fig 2F). The *alms1^−/−^* kidney sections contained apparent interstitial degradation and distal and proximal tubules were dilated compared to control animals (Fig 2F, colored arrows). These observations are consistent with degradation of the tubules and interstitial space and with previously reported phenotypes in *Alms1^−/−^* mice (18).

### *alms1^−/−^* zebrafish exhibit systemic metabolic defects

Having confirmed the presence of previously reported prominent multi-organ features in the *alms1^−/−^* zebrafish (4,18,19), we explored the possibility of metabolic defects. Alström patients exhibit high rates of early onset obesity coupled with hyperinsulinemia and progression to frank T2DM, often as early as the second decade of life (4). To examine the possibility of increased weight gain, we subjected adult wild type or mutant fish clutch-mates to an 8-week dietary regimen consisting of either regular pellet diet (control diet) or a ten-fold increased weight of pellet diet supplemented with five-fold weight of egg yolk powder (high fat diet) (22,23). *alms1^+/+^* fish did not gain significant weight on the control diet, but the high fat diet produced an increase of 1.9-fold over the starting weight at 8 weeks in these animals (p=0.033; Two-Way ANOVA; Fig 3A-B). By comparison, the *alms1^−/−^* fish gained weight under all dietary conditions. Control diet in *alms1^−/−^* fish resulted in a 1.6-fold increase over starting weight and the high fat diet produced a 3.2-fold increase in weight over the 8-week time period. Notably, significant weight gain was apparent after only 3 weeks of high fat feeding (p=0.021; Two-Way ANOVA; Fig 3A-B). We did not observe quantifiable differences in food intake at larval stages, as such we propose that the obesity phenotype is unrelated to central regulation of satiety or hunger (Fig S1B).

**Fig 3.**
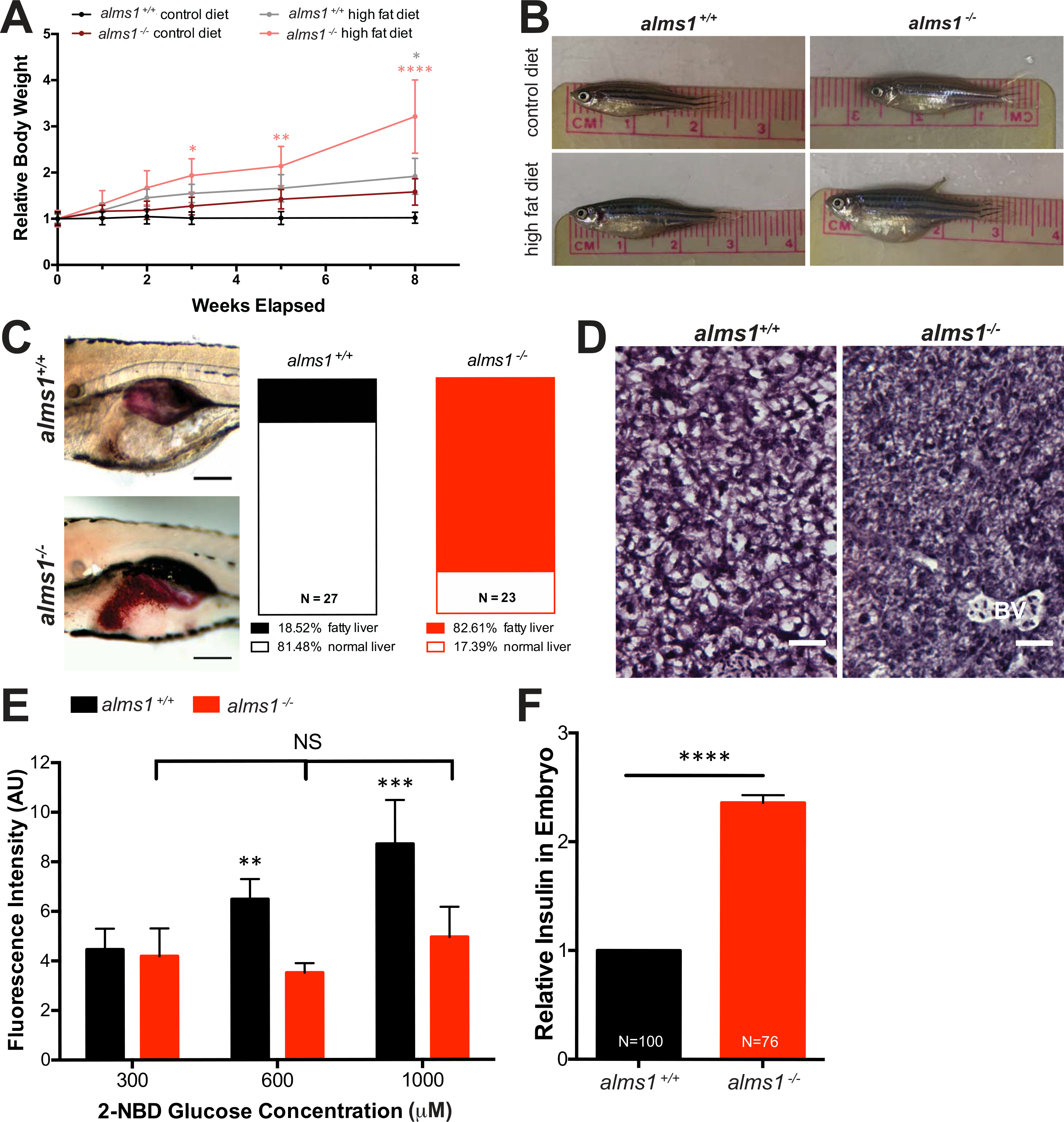
*alms1^−/−^* zebrafish exhibit increased weight gain and systemic metabolic defects. (A) *alms1^+/+^* and *alms1^−/−^* zebrafish at 3 months were fed controlled diets of either maintenance diet (control), or overfeeding with high fat (HF) diet for 8 weeks (n=4-6 animals per condition). Statistics, Two-Way ANOVA compared to *alms1^+/+^* control diet. (B) Representative images of *alms1^+/+^* and *alms1^−/−^* zebrafish after 8 weeks of either control diet or high fat diet. (C) Representative images of Oil Red O staining in livers of *alms1^+/+^* (n=27) and *alms1^−/−^* (n=23) larvae at 6 dpf. Scale bar, 500 μm. Quantification of Oil Red O positive livers. Significance, Chi-squared. (D) Representative regions of sectioned H#E liver tissue from *alms1^+/+^* and *alms1^−/−^* zebrafish at 6 months showing signs of hepatomegaly. Scale bar, 25 μm. BV: blood vessel. (E) FITC-labeled glucose at indicated concentrations was provided to *alms1^+/+^* larvae (n= 5-8 per group) and *alms1^−/−^* larvae (n= 5-12 per group) for 4 hrs. The fluorescence intensity was quantified for each genotype. Statistics, Two-Way ANOVA. (F) Relative insulin by ELISA-based detection in *alms1^+/+^* (n=100) and *alms1^−/−^* (n=76) larvae at 5 days post-fertilization. Statistics, Student’s two-tailed t-test with Welch’s Correction. Where indicated, * p<0.05, **p<0.01, **** p<0.0001.

The truncal obesity typically observed in Alström patients is often accompanied by hepatic steatosis, a common hepatic manifestation of metabolic syndrome. Fatty liver has been reported in a substantial proportion of patients as well as mouse Alström models (4,18). To determine if mutant zebrafish exhibit abnormal accumulation of hepatic lipids, we stained wholemount 6 dpf larvae with Oil Red O and examined neutral lipid content in the liver (Fig 3C). Only 18% of *alms1^+/+^* embryos exhibited observable and dense Oil Red O staining in their livers, while 82% of *alms1^−/−^* embryos exhibited excess fat deposition (p<0.0001; Observed vs Expected; Fig 3C). By adult stages this was accompanied by hypertrophy of hepatopancreata of *alms1^−/−^* fish, demonstrated by increased cell density when compared to age- and section-matched wild type animals (Fig 3D).

Excessive weight gain and hepatic steatosis are indicative of metabolic dysfunction and could suggest insulin resistance and impaired glucose disposal. To evaluate glucose uptake in *alms1* mutant fish, we exposed 6 dpf larvae to 2-(N-(7-Nitrobenz-2oxa-1,3-diazol-4-yl)Amino)-2-Deoxyglucose (2-NBDG), a fluorescently labeled glucose analog, and quantified fluorescence intensity in larval kidneys after 4 hours (24). *alms1^+/+^* animals exhibited a dose-dependent increase in fluorescence upon exposure to increasing concentrations, reaching a 9.75-fold ±3.53 increase over baseline with 1000 μM 2-NBDG (p=0.0003; Two-Way ANOVA; Fig 3E). *alms1^−/−^* larvae, however, did not exhibit any marked increase in fluorescence, reaching only 1.8-fold ± 2.44 over baseline with 1000 μM 2-NBDG (p=0.37; Two-Way ANOVA; Fig 3E), consistent with impaired glucose disposal and, potentially, insulin resistance. To further probe this latter possibility, we quantified insulin levels by high-sensitivity insulin ELISA in pooled protein lysates from 5 dpf larvae and found a 2.36-fold increase in insulin levels in *alms1^−/−^* larvae (p<0.0001; Student’s t-test; Fig 3F), consistent with hyperinsulinemia.

### *alms1^−/−^* zebrafish display impaired β-cell function

While hyperinsulinemia and insulin resistance provide the physiological foundation for diabetes, loss of β-cell mass and function is the hallmark of progression to disease. We previously described increased β-cell apoptosis and decreased β-cell proliferation in *alms1* knockdown in zebrafish larvae (15), indicating a β-cell deficiency in Alström-associated diabetes. Consistent with these data, *alms1^−/−^* adult pancreatic tissue lacked integrity and had fewer insulin granule-positive areas when compared to location-matched control tissues (Fig 4A). At 5 dpf larval stages, *alms1^−/−^* larvae contained an average of 20.4 ± 1.3 β-cells per fish, which is a reduction from the 31.9 ± 1.0 β-cells per fish in controls (p<0.0001; Student’s t-test; Fig 4B). Given the reduced β-cell mass in *alms1^−/−^* animals at all stages, we hypothesized an inability to appropriately sense and regulate systemic glucose in *alms1^−/−^* animals. Proper sensing of high glucose conditions results in expansion of β-cell mass (25). We evaluated this response using *alms1^+/−^* progeny treated with either glucose-free standard embryo media or media supplemented with high glucose (40 mM) from 24-120 hpf. Larvae were fixed and mCherry+ β-cells were imaged and counted prior to genotyping (Fig 4C). This blinded analysis demonstrated β-cell expansion in wildtype siblings grown in 40 mM glucose media from an average of 30.6 ± 1.1 to 36.7 ± 1.9 β-cells per fish (p=0.0049; One-Way ANOVA; Fig 4D), consistent with our previous observations (15). The *alms1^−/−^* larvae again exhibited reduced numbers of β-cells under control conditions, an average of 27.0 ± 1.1 β-cells, and failed to increase the number of β-cells after high glucose exposure, an average of 29.1 ± 0.9 β-cells per fish (p=0.0671; One-Way ANOVA; Fig 4E), implicating Alms1 in the β-cell response to elevated systemic glucose.

**Fig 4.**
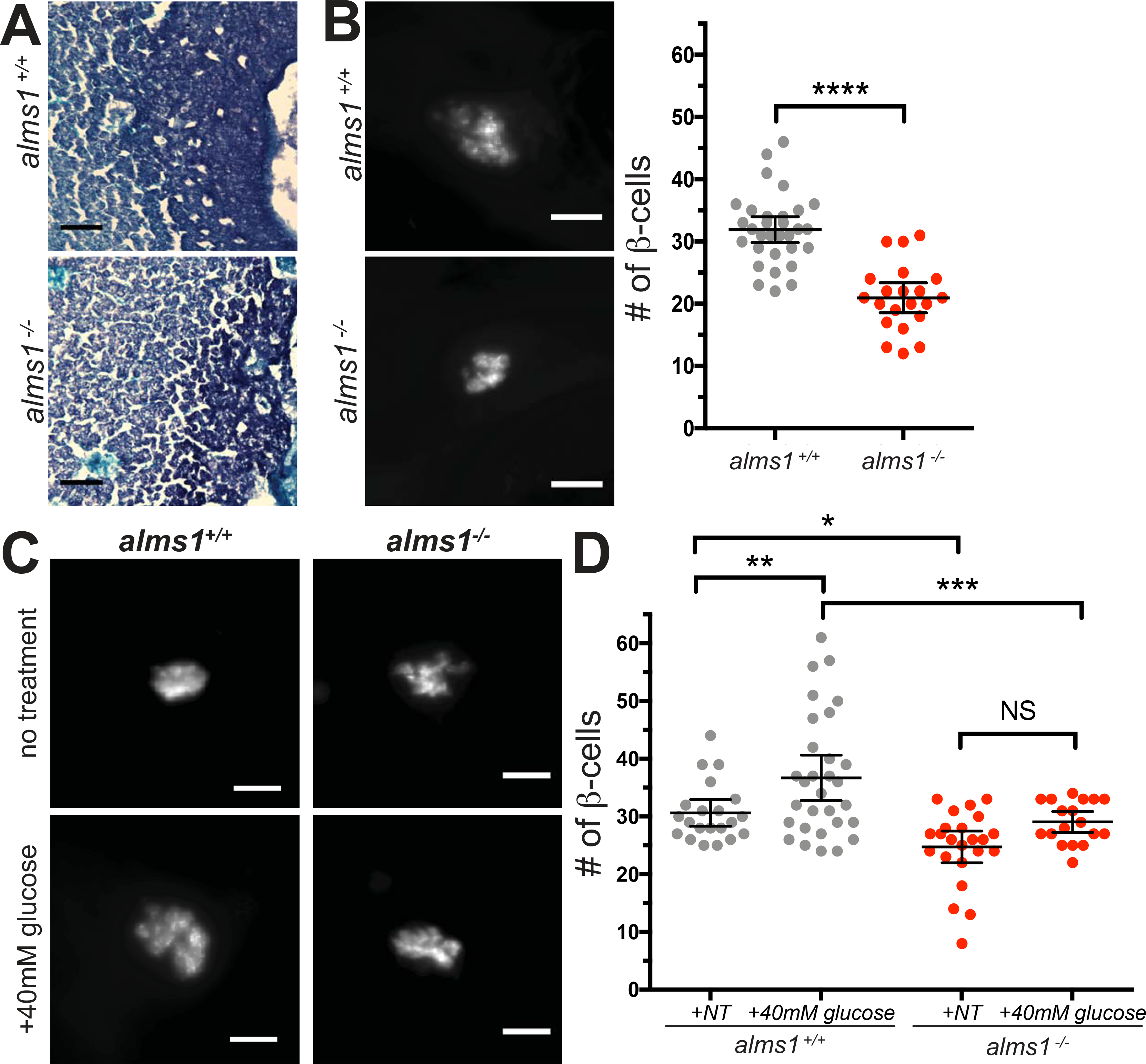
*alms1^−/−^* zebrafish islets show reduced size and glucose responsiveness. (A) Aldehyde fuchsin staining of sectioned zebrafish at 6 months showing aberrant islet structure (dark purple regions) in *alms1^−/−^* as compared to *alms1^+/+^*. Scale bar, 50 μm. (B) β-cell imaging and quantification of 5 dpf larvae in *alms1^+/+^* (n=31) and *alms1^−/−^* (n=22) larvae. Scale bar, 25 μm. Statistics, Student’s t-test with Welch’s Correction. (C) Representative images of β-cells with and without exposure to 40 mM glucose in *alms1^+/+^* and *alms1^−/−^* larvae. Scale bar, 25 μm. (D) Quantification of β-cells at 5 dpf from *alms1^+/+^* (NT=22, Glu=31) and *alms1^−/−^* (n, NT=25, Glu=18) larvae. Statistics, One-Way ANOVA. Where indicated, * p<0.05, ** p<0.01, *** p<0.001, **** p<0.0001.

The β-cell deficits and elevated systemic insulin suggest a pathophysiology in which β-cells are hypersecretory and also unable to properly regulate insulin secretion in response to exogenous cues. To better understand how loss of *alms1* may impact β-cell function, we generated single-cell homogenates from 5 dpf larvae of either wildtype Tg(*insa:mCherry*) or Tg(*insa:mCherry*)*;alms1^−/−^* and isolated the mCherry+ cells via FACS sorting (Fig 5A) (26). Using RNA from the isolated cells, we carried out whole transcriptome analysis via RNA-Seq (26,27). Using a cut-off for differentially expressed genes of a fold change of greater than 1 or less than – 1, we identified a total of 3,880 up-regulated genes and 5,531 down-regulated genes in the *alms1^−/−^* β-cells compared to β-cells isolated from wild type larvae (Fig 5B). We first confirmed the β-cell-enriched nature of these cells by confirming elevation of islet and β-cell markers, relative to markers of other cell types (Fig 5C). We then carried out pathway analysis of significantly changed genes and found that the down-regulated genes in mutant β-cells strongly supported a generalized decrease in cellular activities, protein and RNA processing, consistent with the significant apoptosis that we previously reported (Fig S2) (15). Over-represented terms in the set of upregulated genes were in large part related to cellular transport and secretion (37 of 74 categories) and cell membrane ion transport (Fig S3A). These findings support a role for *alms1* in regulation of secretion and membrane depolarization in β-cells. We also identified a group of significantly abundant pathways in the up-regulated genes with known pancreatic functions (Fig S3B). These genes can be clustered into those implicated in monogenic diabetes, nutrient transport (mainly metal ion and amino acids carriers), insulin secretion (including calcium and potassium channels), and generalized pancreatic secretion (Fig 5C). The intersections of these pancreas-related pathways contain a subset of six genes, including slc2a2/glut2, found in 3 pathways, and insulin, found in 2 pathways (red numbers, Fig 5C).

**Fig 5.**
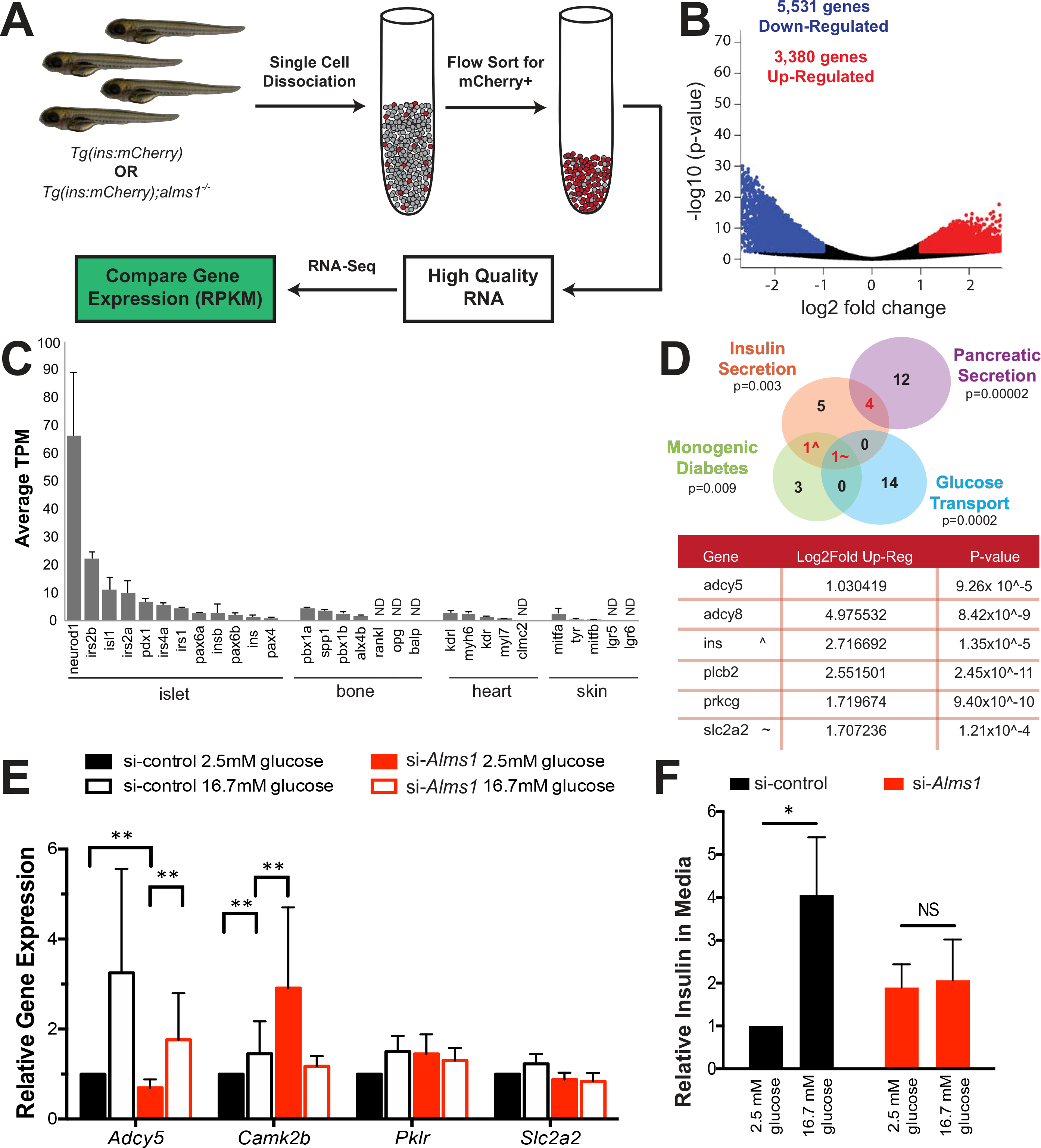
Alms1 influences glucose sensing in β-cells. (A) Schematic of experimental design for comparative gene expression in β-cell enriched populations from age-matched control and *alms1^−/−^* larvae. (B) Volcano plot of significantly differentially expressed genes from GRZ10 between control and *alms1^−/−^* larvae. (C) Selected tissue specific genes identified in isolated *alms1^+/+^* cells. Note the presence and high expression of pancreatic markers. ND: not identified. (D) Subset of significantly up-regulated pathway nodes, identified by ConsensusPath DB, in *alms1^−/−^* β-cells. Genes within intersections are listed in table alongside the fold-increase and significance compared to *alms1^+/+^* β-cells. (E) Expression of glucose response genes in si-*Alms1* β-cells under basal (2.5 mM) and high glucose (16.7 mM) conditions (n=4). Cells were collected after 10 minutes after glucose stimulation. Statistics, Two-Way ANOVA. (F) Relative insulin by ELISA-based detection in culture media from β-cells after 30 minutes of exposure to 2.5 mM and 16.7 mM glucose (n=3). Normalized to basal si-control. Note failure of si-*Alms1* cells to alter insulin secretion. Statistics, Two-Way ANOVA. Where indicated, * p<0.05, ** p<0.01.

### Depletion of *Alms1* in cultured β-cells impairs glucose challenge response

Given the identified functional defects in *alms1^−/−^* β-cells and the implication of insulin secretion defects from transcriptomic data, we hypothesized that the diabetes phenotype may be linked to dysregulated insulin secretion in the absence of Alms1. Thus, we evaluated the mechanism by which ALMS1 impacts insulin secretion using a simplified cultured mouse β-cell model, MIN6.

*Alms1* knockdown was accomplished via transient siRNA transfection (si-*Alms1*) with an average of 0.38 ± 0.04 *Alms1* expression compared to control levels (p<0.0001; Student’s t-test; Fig S1C). We next examined the genes implicated by RNA-Seq in cultured β-cells. When exposed to 16.7 mM glucose, physiological high glucose conditions, control cultured β-cells showed an increase in adenylyl cyclase 5 (*Adcy5*) and calcium/calmodulin-dependent protein kinase 2b (*Camk2b*), both known to transmit signals downstream of glucose transport in murine β-cells, along with moderate changes to *Pklr*, which initiates intracellular glucose processing and alters cellular metabolism, and *Slc2a2*, the main murine glucose transporter (Fig 5E) (28). Of note, the si-*Alms1* β-cells exhibited a dampened *Adcy5* response and higher levels of *Camk2b* gene expression profiles (p=0.002; Two-Way ANOVA) irrespective of glucose levels (Fig 5E). This appeared to be accompanied by a failure to alter *Pklr* and *Slc2a2* expression in response to glucose (Fig 5E). In fact, *Pklr* appeared to mimic high glucose control conditions even in si-*Alms1* β-cells cultured in low glucose conditions (Fig 5E), suggesting an inability to properly sense glucose and regulate gene expression.

These gene expression data led us to hypothesize that the si-*Alms1* β-cells would potentially exhibit inappropriate insulin secretion in response to high glucose. The secreted insulin upon stimulation with physiological high glucose in si-*Alms1* β-cells was evaluated using high-sensitivity ELISA based quantification. The control β-cells required 10 minutes to begin secreting insulin in response to high glucose, and reached 2.2-fold the unstimulated insulin levels by 30 minutes (Fig 5F). The si-*Alms1* β-cells showed 1.6-fold higher levels of secreted insulin than control β-cells without glucose stimulation (Fig 5F). After 30 minutes of high glucose, the si-*Alms1* β-cells were at only 0.88-fold of unstimulated control levels (p=0.032; Two-Way ANOVA; Fig 5F). These data suggest a modest hypersecretory basal state in unstimulated Alström β-cells accompanied by an impairment in glucose-stimulated insulin secretion.

## DISCUSSION

These studies characterized the physiologic and phenotypic responses of a newly generated Alström model created in zebrafish using CRISPR/Cas9. This new mutant line not only exhibits reduced Alms1 RNA and protein levels with body dysmorphogenesis of known zebrafish ciliary genes (Fig 1) (17,29), but demonstrates the known physiological defects of both Alström patients and the mouse Alström model, including functional retinal degradation and renal tubule dilation (Fig 2) (4,18). Given the molecular changes and phenotypic differences, we concluded this Cas9-induced deletion represents a loss-of-function mutant model of the obesity ciliopathy, Alström syndrome.

Since a defining characteristic of this ciliopathy is obesity, we examined and demonstrated a propensity for excess weight gain and inappropriate lipid storage, implicating Alms1 in zebrafish metabolic regulation (Fig 3). These conserved effects are of particular interest as the metabolic organs of the zebrafish closely mimic those of human physiology (13,14). In particular, adipose depot locations, and governing hormones are all conserved between zebrafish and humans (30,31). Furthermore, as is often found in the general obese population, Alström patients exhibit hyperinsulinemia, insulin resistance and type 2 diabetes mellitus (32). The *alms1* mutant zebrafish displayed characteristics supporting a conserved role for the gene. Namely, the *alms1^−/−^* fish are hyperinsulinemic, fail to respond to glucose challenges and have islets with reduced β-cell numbers (Fig 3-4). Beyond the reduced overall β-cell mass, the remaining β-cells in *alms1^−/−^* animals are unhealthy and improperly functioning. RNA-Seq gene expression analysis in cells isolated from *alms1^−/−^* animals show excess secretory activity, especially in insulin-related genes, and cellular transport alongside an overall decrease in transcription and translation (Fig 5, Fig S2-S3). Consistent with this conclusion was the hyperinsulinemia in larval zebrafish and insulin hypersecretion in unstimulated cultured β-cells. Glucose-stimulated insulin secretion was, however, severely impaired in cultured β-cells as was the response of larval β-cells to high glucose conditions. These defects combined with the significant changes in expression of genes involved in secretory pathways lead us to conclude that unstimulated β-cells without *alms1* are hypersecretory.

Based on these observations, we propose unstimulated basal β-cell hypersecretion as a primary driver of hyperinsulinemia in *alms1^−/−^* animals. Insight from RNA-Seq gene expression changes and observed phenotypic responses lead us to propose two aspects of β-cell function that are reliant upon Alms1: 1) control of insulin secretion in the absence of stimuli and 2) appropriate β-cell glucose sensing. We used a simplified β-cell culture system to examine these aspects individually. Knockdown of *Alms1* resulted in excess secreted insulin without glucose stimulation, but failure to respond to high glucose stimulation at the level of either insulin secretion or altered gene expression (Fig 5). Epistatic analysis of the glucose-sensing and insulin response pathways in *Alms1* knockdown or knockout β-cells will be important to identify the exact genetic pathway defects in those cells and could be confirmed in the *alms1^−/−^* animals. The molecular mechanism by which Alms1 impacts secretion is unknown, and such studies would inform the function of the Alström protein in β-cells and potentially provide insight into its role in other cell types. Importantly, our observations provide additional evidence of a central role for β-cell cilia in sensing of extracellular stimuli that directly impact insulin secretion. Ciliary localization of the insulin receptor in β-cells, for example, is critical to normal function (33).

Several adenylyl cyclases, including those encoded by *ADCY5* and *ADCY8*, have also been linked to cilia in other cell types (34,35) and are critical for cAMP generation, a major regulator of glucose-stimulated insulin secretion (36). It is therefore possible that disruption of Alms1 impacts the ability to sense systemic insulin or glucose levels, resulting in a dysregulation of β-cell response to stimuli and subsequent secretion.

While the early onset insulin hypersecretion may contribute to systemic insulin resistance and even lead to eventual β-cell exhaustion, this does not rule out the implication of peripheral tissue insulin resistance that is also a major trigger of T2DM pathology. Given the non-tissue specific nature of this mutant, the individual contributions of adipose or muscle tissue to the observed diabetes phenotypes cannot be evaluated. The primary role of β-cell dysfunction in T2DM versus a secondary defect to peripheral insulin resistance is currently a topic of debate (37,38). Generation of tissue-specific deletion of *alms1*, including peripheral tissues and β-cell-specific knockout, will permit analysis of the cell-autonomous role for Alms1 in the context of systemic dysfunction. The utility of the zebrafish and its recapitulation of human disease phenotypes with high fidelity suggest that these genetic models of diabetes can be useful tools in isolating the root cause of β-cell failure and T2DM etiology.

## MATERIALS AND METHODS

### Zebrafish Husbandry and Stocks

Experiments were performed using the Tg(*insa:*mCherry) from the Zebrafish International Resource Center. Adult fish were screened and a 7 bp deletion in exon 4 of *alms1* was identified, resulting in the Tg(*insa*:mCherry); *alms1^−/−^* line. Embryos were raised at 28.5 °C for all experimental analyses. Adult zebrafish were housed and mated naturally, based upon standard protocols (ZIRC). All zebrafish work was conducted in accordance with the University of Maryland Baltimore IUCAC guidelines.

### Adult Zebrafish Analyses and Imaging

For histological analyses, adult fish—approximately six months in age—were sacrificed in 0.05% tricaine (3-amino benzoic acid ethyl ester) until unresponsive (Sigma). The tail segment behind the anal fin was removed using a razor blade and samples were fixed for 24 hrs in Dietrich’s solution (30% ethanol, 10% formalin, 2% glacial acetic acid) then decalcified in 0.5 M ethylenediaminetetraacetic acid for 7 days. Samples were processed for cryo-sectioning using serial sucrose gradient before being embedded in optimal cutting temperature (OCT) compound (Tissue-Tek). Samples were sectioned into 10 μM thick transverse sections and mounted. Hemotoxylin (Thermo Scientific) and Eosin Y (95%, Sigma) staining was performed to evaluate overall tissue morphology. Aldehyde fuchsin staining (18) with Fast Green counterstain (Sigma) was performed to examine insulin+ tissues. Images were collected on a Nikon Eclipse at 10-40X.

Visual responses were examined in adult fish—approximately nine months old—that were dark adapted overnight. All following procedures were done under dim red light. Fish were anesthetized by submersion in tricaine solution. After anesthesia, zebrafish were transferred to wet Whatman filter paper stack and immobilized by injection of 10 μL of 25 mg/ml gallamine triethiodide (Enzo Life Sciences) below the gill. Anesthesia was maintained with continuous perfusion of tricaine-containing oxygenated system water. The reference AgCl electrode was placed near the eye and the recording electrode was placed on the eye. The fish with attached electrodes was placed in a Ganzfeld chamber and presented with increasing scotopic stimuli (0.025-9.952 cd x s/m2) at 10-60 sec intervals using UTAS BigShot (LKC Technologies). A minimum 3 waveforms per intensity were averaged.

Adult feed studies were carried out in 1 L tanks containing genotyped clutch-mates starting at 3 months of age. The volumes of food were modified from published protocols (22,23). Control diet volume was assessed prior to beginning the experiment as the amount of food consumed by wild type fish in 5 minutes. Fish were fed once a day with assigned dietary conditions and the length and weight were quantified at indicated time points. The food volume was calculated after each weight measurement to account for weight gain throughout the study period.

All image processing and analyses were carried out using Fiji (39).

### Embryonic Zebrafish Analyses and Imaging

Fat content in larval livers were examined after yolk absorption was complete, 5-7 dpf, via Oil Red O staining of whole animals fixed in 4% paraformaldehyde (PFA). Fatty liver was identified by Oil Red O accumulation in the liver area, found on the right side of yolk-depleted larvae. All images were collected on a Zeiss Axioscope.

β-cell analyses were performed at the indicated time points using previously published protocols (15,25,40). Briefly, the mCherry+ cells were counted β-cells in larvae containing the Tg(*insa:mCherry*) transgene after fixation in 4% PFA and post-fixation in methanol. Larvae were placed with the left lateral side facing the slide and compressed under coverslips with Prolong Gold antifade (Life Technologies) media. Tg(*insa:mCherry*);*alms1^+/−^* progeny were glucose challenged using medium supplemented with 40 mM glucose at 24 hpf and collected at 5 dpf for β-cell analysis. Control conditions were standard embryo medium. After β-cell counts were collected the *alms1* genomic identity of each larva was determined via PCR genotyping. Images were collected on a Nixon W1 using a 40x objective.

Embryonic feeding assays were performed as previously published (41). Briefly, equal amounts of BODIPY FL C12 (Life Technologies) egg yolk were provided to 7 days post fertilization (dpf) larvae for a period of 4 hours at 28.5 °C. The embryos were then lysed and total fluorescence at 510 nm was quantified.

Glucose uptake and insulin levels were evaluated in 5 dpf larvae. 2-(N-(7-Nitrobenz-2oxa-1,3-diazol-4-yl)Amino)-2-Deoxyglucose (2-NBDG) in dimethyl sulfoxide (DMSO) was provided to larvae at 300-1,000 μM in embryo medium at 28.5 °C for 4 hrs. The larvae were then rinsed 4X with embryo medium and anesthetized before imaging. All images were collected on a Zeiss Axioscope. Glucose uptake was evaluated as total fluorescence in the embryonic kidney by ImageJ, relative to DMSO alone. Larval insulin levels were evaluated in pooled larval lysates, mechanically dissociated in Nonidet P-40 protein buffer, using high-sensitivity insulin ELISA (Mercordia).

All image processing and analyses were carried out using Fiji (39).

### RNA-sequencing of Isolated Preparation β-cells

5 dpf larvae from Tg(*insa:mCherry*) and Tg(*insa:mCherry*);*alms1^−/−^* were dissociated into single cells and sorted via mCherry+ signal using a BD FACS Aria II (BD BioSciences) (26). RNA was extracted from isolated cell fraction using an RNA extraction kit (Qiagen). RNA quantity and quality were assessed via 260/280 absorption. Samples were provided in duplicate for library preparation and quantitative analysis using Next Generation Sequencing and an Illumina HiSeq 2×150 PE (GENEWIZ). Fragments were aligned to the GRZ10 genome with CLC Genomics Server program v10.0.1. Full expression datasets are deposited at GEO and available upon request.

### Analyses of RNA-Seq Expression datasets

Analysis of the identified list of significantly differentially expressed genes was performed as previously published (26). Pathway interactions were visualized using ConsensusPath DB (42,43) and ontology clusters were evaluated using the Gene Ontology Consortium (44,45).

### Cell Culture

Culture of MIN6 cells (CRL11506; American Type Culture Collection) were cultured in DMEM-H (American Type Culture Collection) supplemented with 15% heat-inactivated fetal bovine serum (FBS) and 1X penicillin/streptomycin (Sigma). Knockdowns were accomplished using Lipofectamine 3000 (Life Technologies) and either scrambled control or *Alms1*-targeted siRNA (Life Technologies). Efficacy of siRNA knockdown was evaluated via qRT-PCR.

Glucose stimulation of cultured β-cells was performed on cells plated at equal densities, as determined by hemocytometer counts, using 2.5 mM and 16.7 mM glucose as baseline and high glucose concentrations, respectively. Insulin was assessed in media collected at the indicated time points using high-sensitivity insulin ELISA (Mercordia).

### Western blots

5 dpf zebrafish were mechanically homogenized using mortar and pestle in NP-40 buffer supplemented with protease and phosphatase inhibitor (Sigma). Homogenized tissue was incubated on ice for 15 mins with regular vortexing to lyse. Lysates were centrifuged at 9,400 g for 10 minutes and the supernatant was collected. Lysate supernatant was boiled with Laemmli sample buffer at 95 °C for 10 mins. Equal amounts of protein were loaded and transferred onto a nitrocellulose membrane. Membranes were blocked for 1 hr in Tris-buffered saline (TBS) with 0.1% Tween-20 and 5% BSA. Membranes were then incubated overnight with Goat Anti-Alms1 (1:500; Abcam) or Rabbit Anti-Actin (1:2,000; Sigma). Detection of specific proteins was accomplished after 1 hr incubation with species specific HRP-conjugated secondary antibody (1:15,000; Jackson Immuno) and application of ECL Substrate (Pierce). Protein intensity was normalized to actin and quantified via densitometry function in ImageJ.

### qRT-PCR

RNA was extracted using RiboZOL RNA Extraction Reagent (VWR) and converted to cDNA via the FirstStrand cDNA Synthesis kit (Thermo Scientific), according to manufacturer protocols. Target gene expression was determined on a LightCycler 480 (Roche) using 2X SYBR Green Master Mix (Roche) and compared by ΔΔCT. Actin and GADPH were used as control in zebrafish and cultured cells, respectively. Primer sequences are available upon request.

### Statistical Analysis

All experiments represent a minimum of three replicates, with the number of analyzed samples (n) provided. Prism 6.0 software (GraphPad) was used to determine appropriate analyses and statistical significance and is provided in individual figures.

### Ethics Statement

All animal research was approved by the University of Maryland Baltimore Institutional Animal Care and Use Committee (IACUC), approval #0116027.

## AUTHOR CONTRIBUTIONS

JEN designed and performed experiments, analyzed data, and wrote the manuscript. TLH generated the mutant line and performed the dietary experiments. CCL perform Oil Red O experiments. RM performed embryonic dietary assays. SL assisted in mutant line generation. MSM and RH performed cryopreservation and tissue sectioning. SS and ZMA performed photoreceptor function tests. CJW contributed the FACS sorting experiment. NAZ contributed experimental design and wrote the manuscript.

## ACKNOWLEDGEMENTS

The authors wish to thank the University of Maryland School of Medicine Center for Innovative Biomedical Resources, Confocal Microscopy Core in Baltimore, MD for equipment use and assistance with image acquisition. Jonathan Van Ryzin and Margaret McCarthy’s laboratory at the University of Maryland, Baltimore were invaluable in the acquisition of the histology images. The zebrafish β-cell isolation was carried out with assistance from and use of the CCR Frederick Flow Cytometry Core.

This work was funded by the grants from the National Institutes of Health in Bethesda, MD. Specific funding was provided by grants from the National Institutes of Diabetes, Digestive and Kidney Disorders—R01DK102001 (NAZ), P30DK072488 (NAZ and RM), T32DK098107 (JN and TLH), and F31DK115179 (TLH)—and the National Institute on Deafness and other Communication Disorders—R01DC013817 (RH), R01DC016295 (ZMA), and F31DC016218 (MSM).

---

**Fig S1. Organ defects in *alms1^−/−^* zebrafish.**

(A) Zebrafish *alms1* gene expression in adult fish of indicated genotype. Statistics, One-Way ANOVA. (B) Total fat consumed as assessed by fluorescence per 20 larvae in *alms1^+/+^* (n=6) and *alms1^−/−^* (n=7) larvae at 7 dpf. (C) cDNA levels of Alms1 in cultured MIN6 β-cells in control and si-*Alms1* conditions. Statistics, two-tailed Student’s t-test with Welch’s Correction. Where indicated, **** p<0.0001, NS not significant.

**Fig S2. Significantly down-regulated genes in *alms1^−/−^* β-cell enriched populations.**

Gene Ontology (A) and ConsensusPath DB (B) analysis of significantly down-regulated genes between β-cells isolated from *alms1^+/+^* and *alms1^−/−^* zebrafish. Green highlight, incretin hormone pathway nodes.

**Fig S3. Significantly up-regulated genes in *alms1^−/−^* β-cell enriched populations.**

Gene Ontology (A) and ConsensusPath DB (B) analysis of significantly up-regulated genes between β-cells isolated from *alms1^+/+^* and *alms1^−/−^* zebrafish. Green highlight, pancreatic secretion and diabetes-related pathway nodes.

